# Re-evaluating the benefit of training scale in cell foundation models

**DOI:** 10.64898/2026.06.04.730272

**Authors:** Haroon M Shakeel, Mengyuan Shen, Stefano Mangiola

## Abstract

Single-cell foundation models are self-supervised neural networks pretrained on millions of cellular transcriptomes to learn representations that transfer across biological domains and downstream tasks. Despite increasing investments in model and pretraining data scale, it remains unclear how to assess whether these investments improve the quality of learned biological representations. Recent evidence from standard finetuning benchmarks suggests that cell foundation models reach performance saturation after exposure to only a fraction of the available pretraining data. We hypothesised that conventional finetuning may obscure improvements in underlying representation quality and the biological information they encoded during pretraining. To address this, we tested second- and first-generation cell foundation models across corpus sizes using standard finetuning and extended it with a comprehensive fully frozen evaluation. We found that the training-data scaling relationship becomes apparent under fully frozen finetuning, particularly for second-generation models. Zero-shot analyses similarly recovered dependence on pretraining scale. Our findings reveal a fundamental limitation of conventional foundation model evaluation and establish the importance of frozen-representation benchmarking for assessing representation quality and guiding pretraining-scale decisions.

## Main

Deep neural networks have become a major framework for learning representations from high-dimensional data ^1,2^. In natural language processing, transformer-based models pretrained on large collections of unlabelled text have demonstrated emergent intelligence that is transferable across domains and tasks ^3–5^. Single-cell genomics has increasingly adopted this paradigm, developing cell foundation models on large single-cell atlases ^6–15^. These models promise to generalise to downstream tasks such as cell type annotation, perturbation response prediction and state imputation. The central assumption is that exposure to a large and diverse cell compendium enables models to learn reusable representations across cell types, tissues, demographics, disease states, and developmental stages. Consequently, progress in the field is often framed in terms of scale, with larger models and larger pretraining corpora representing key methodological advances ^8,10,14,15^.

A fundamental challenge in foundation model development is determining whether increasing pretraining scale improves the quality and transferability of learned representations. Standard finetuning benchmarks provide limited insight into this question because downstream performance reflects both the quality of the pretrained representation and the adaptive capacity of unfrozen layers. Indeed, recent benchmarks have reported early performance saturation despite substantial increases in pretraining data^16^. Whether this saturation reflects limitations in representation learning or the evaluation strategy itself remains unclear.

Here, we systematically investigated how pretraining scale, model capacity and evaluation strategy jointly influence the measured performance of cell foundation models. By comparing partially unfrozen and fully frozen evaluation regimes, we distinguished task-specific adaptation from the contribution of pretrained biological representations. We hypothesised that standard finetuning, where the last encoder layers are unfrozen, can mask genuine foundation representation qualities, and therefore pretraining-scale effects, by allowing a weaker representation to be compensated during task adaptation. That is, the dependence of performance on the pretraining scale, useful to assess the benefit of large corpora, would emerge only when downstream prediction relies solely on the foundation representation. To test this, we pretrained second- and first-generation Geneformer variants ^8,15^ (comprising 104 million and 10 million parameters, respectively) across balanced and random corpus-reduction series spanning 34 million to 266k cells, and evaluated each model under partially unfrozen and fully frozen finetuning regimes. We further assessed whether these models could recover functional gene networks and predict perturbation responses in zero-shot settings.

### A curated cell compendium allows a reproducible, high-quality benchmark

To improve the quality and reproducibility of our benchmark, we developed a data subsampling pipeline and API for Python and R (cellnexus.org/subsampling) based on the curated cell compendium cellNexus ^17^ (Figure 1A). This pipeline and resource offer a quality-controlled and reproducible platform for cell foundation model evaluation, including high-quality data sources and subsampling routines. Using balanced and random subsampling strategies, we progressively reduced the pretraining data from 34M to 266k cells, resulting in a total of 19 corpora. In this study, balancing was applied across samples and cell types, which preserved proportionally more cells from smaller samples and rare cell types (Figure 1A, B; Supplementary Figure S1A, B). Using these corpora, we pretrained the second- and first-generation Geneformer variants from scratch ^8,15^, totalling 38 foundation models pretrained for 12,104 GPU-hours. As expected, progressive subsampling led to a linear reduction in training time (Figure 1C). Relative to 34M cells, pretraining on 266k cells reduced training time by approximately 93-fold for the second-generation model and 200-fold for the first-generation variant, falling from 801 to 8.6 hours and 204 to 1.02 hours, respectively.

**Figure 1.**
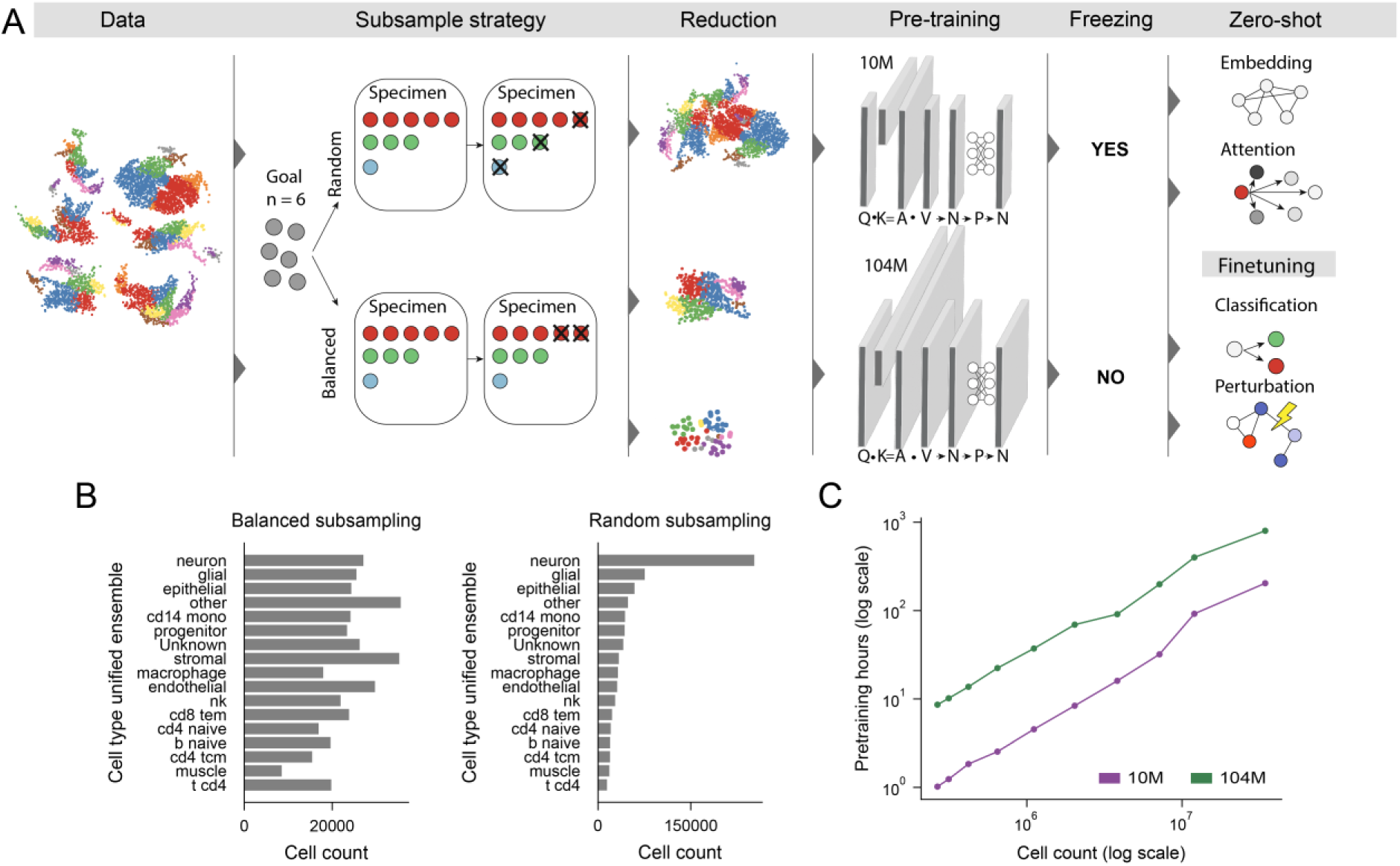
Overview of the corpus subsampling, pretraining, and evaluation framework. **A**, Schematic of the study design, including corpus subsampling, pretraining of the second- and first-generation Geneformer variants, and downstream evaluation under fully frozen and partially unfrozen regimes. **B**, Distribution of the 17 most abundant cell types after random subsampling of 1 million cells from cellNexus, compared with their corresponding distributions after cell-type-balanced subsampling. **C**, Pretraining time as a function of corpus size for the second-generation (green) and first-generation (purple) Geneformer variants, shown on log-scaled axes.

We benchmarked each pretrained model across five common finetuning tasks spanning task-specific datasets under the standard partially unfrozen and fully frozen finetuning regimes (see Methods). This produced 380 task-adapted models. We additionally finetuned the original Geneformer models pretrained on Genecorpus and released with their respective studies ^8,15^, adding 20 further task-adapted models and bringing the total to 400. Partially unfrozen finetuning updated the prediction head, input embedding layer, and selected upper encoder layers, whereas fully frozen finetuning updated only the prediction head. This comparison examined whether similar performance after encoder adaptation concealed differences in the task-relevant information accessible from fixed foundation representations. We also evaluated HLA-module recovery and attention-derived regulatory associations as complementary biological probes that required no task-specific parameter updates and assessed the biological information encoded in the foundation representations (see Methods).

### Standard finetuning early performance saturation extends to second-generation models

To determine whether the previously reported early downstream performance saturation ^16^ extends to second-generation cell foundation models, we compared Geneformer variants as a representative generational increase in model capacity. We compared the second-generation (104 million parameters) with first-generation (10 million parameters) Geneformer variants under standard, partially unfrozen finetuning. Overall, in these settings, our results matched the known saturation of early finetuning performance for the second- and first-generation variants.

We first evaluated cell-type classification, a task widely used to benchmark single-cell foundation models ^8,10,14^ (Figure 2A). Under balanced subsampling, the second-generation variant showed a statistically supported but modest positive association with pretraining corpus size (FDR = 0.0020; slope = 0.018), with most improvement confined to smaller corpora. Mean macro-F1 increased from 0.934 ± 0.002 at 266k cells to 0.964 ± 0.0007 at 3.8M cells, compared with 0.966 ± 0.0002 at the full 34M-cell corpus. Despite this association, the predefined saturation threshold was already reached at 266k cells, corresponding to approximately 0.78% of the full cellNexus corpus. The first-generation variant showed even weaker dependence on corpus size (FDR = 0.0037; slope = 0.0052), with mean macro-F1 changing only from 0.949 ± 0.001 at 266k cells to 0.958 ± 0.001 at 34M cells and saturation again occurring at 266k cells. The relative performance of the two variants depended on corpus size: the second-generation variant underperformed up to 1.1M cells and exceeded the first-generation variant only moderately at larger scales. Random subsampling produced the same early saturation point for both variants and similarly showed a weak dependence on pretraining corpus size (Supplementary Table 1), indicating that the observed trend was not specific to the balanced subsampling procedure.

**Figure 2.**
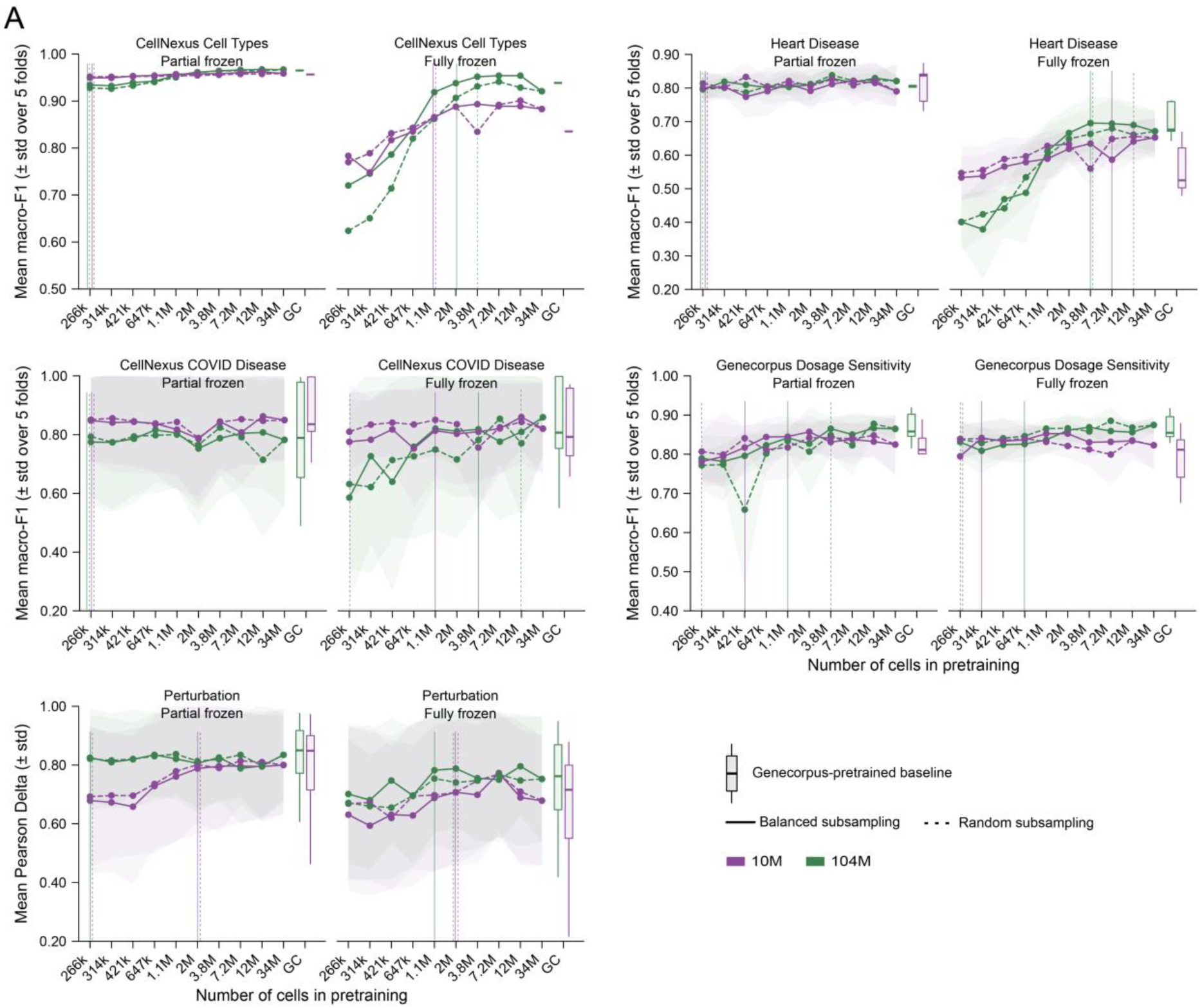
Effects of pretraining corpus size and finetuning regime on downstream performance. **A**, Finetuning performance of second- and first-generation Geneformer variants under fully frozen and partially unfrozen regimes across five finetuning tasks. Performance is shown as macro-F1 score across five-fold cross-validation for classification tasks and Pearson delta for the perturbation prediction task. Vertical lines show the saturation point for the respective model and subsampling strategy.

A similarly limited dependence on pretraining corpus size was observed for disease-status classification. For heart-disease prediction ^8^, the second-generation variant showed no significant association under balanced subsampling (FDR = 0.059), with mean macro-F1 increasing only from 0.796 ± 0.03 at 266k cells to 0.820 ± 0.04 at 34M cells. The first-generation variant was likewise insensitive to corpus size (FDR = 0.44), changing mean macro-F1 from 0.799 ± 0.04 to 0.790 ± 0.04 across the same range. COVID-19 prediction followed the same pattern: performance was not significantly associated with corpus size for either the second-generation variant (FDR = 0.69; mean macro-F1 0.774 ± 0.21 at 266k cells and 0.782 ± 0.22 at 34M cells) or the first-generation variant (FDR = 0.93; mean macro-F1 0.847 ± 0.14 and 0.849 ± 0.13, respectively). All four variant–task combinations reached saturation at 266k cells. Random subsampling reproduced these saturation points for both disease-classification tasks, showing that the absence of a meaningful pretraining-scale effect was robust to the subsampling strategy. Notably, the second-generation variant performed slightly worse than its smaller predecessor on COVID-19 prediction, further indicating that increased model size did not confer a consistent advantage on these narrow disease-classification tasks under standard, partially unfrozen finetuning.

Drug dosage-sensitivity classification ^8^ showed a modestly stronger, although still rapidly saturating, dependence on pretraining corpus size. Under balanced subsampling, the second-generation variant showed a significant positive association with corpus size (FDR = 0.0036; slope = 0.040), with mean macro-F1 increasing from 0.789 ± 0.05 at 266k cells to 0.865 ± 0.03 at 3.8M cells and remaining comparable at 34M cells (0.864 ± 0.02). Performance saturation was reached at 1.1M cells. The first-generation variant showed no significant overall association (FDR = 0.25), although performance initially increased from 0.780 ± 0.07 at 266k cells to 0.844 ± 0.03 at 647k cells before plateauing and reaching 0.824 ± 0.07 at 34M cells. Its saturation threshold occurred at 421k cells, earlier than for the second-generation variant. As observed for cell-type classification, the relative advantage of the larger second-generation variant depended on data availability: it performed worse at corpus sizes up to 2M cells but consistently outperformed the first-generation variant at larger scales. For the second-generation variant, random subsampling delayed saturation from 1.1M to 3.8M cells and modestly reduced performance at the smaller corpus sizes. For the first-generation variant, random subsampling instead shifted saturation from 421k to 266k cells.

Beyond classification, we evaluated perturbation-response prediction using the Norman double-perturbation benchmark ^18^ (Figure 2A). We prioritised Pearson delta over mean *L*_2_ distance because it captures the direction of perturbation-induced transcriptional change and is less sensitive to systematic under- or over-estimation of response magnitude ^19^; mean *L*_2_ distance is reported in Supplementary Table 2. Under balanced subsampling, the second-generation variant showed no significant association between mean Pearson delta and pretraining corpus size (0.824 ± 0.14 at 266k cells and 0.834 ± 0.15 at 34M cells; FDR = 0.68) and reached saturation at 266k cells. The first-generation variant showed a stronger positive association up to 3.8M cells, increasing mean Pearson delta from 0.679 ± 0.25 at 266k cells to 0.790 ± 0.20 at 3.8M cells (FDR = 0.0037; slope = 0.070), after which performance plateaued; saturation was reached at 2M cells. The second-generation variant substantially outperformed the first-generation variant below 2M pretraining cells, but this difference narrowed at larger corpus sizes. Random subsampling produced comparable trajectories for the second-generation variant, with no significant difference in pretraining-scale dependence between the two subsampling strategies (FDR = 0.98; slope = 0.00032), while for the first-generation variant the association was identical to that of balanced subsampling (FDR = 0.0037; slope = 0.061). The saturation points were unchanged at 266k cells for the second-generation variant and 2M cells for the first-generation variant. Thus, although the subsampling strategy slightly affected absolute performance in selected classification settings, it did not materially alter the broader pattern of early performance saturation under partially unfrozen finetuning.

Across both Geneformer variants, standard partially unfrozen finetuning showed early performance saturation under balanced and random subsampling. This pattern is consistent with previous observations of downstream performance plateaus^16^. However, these findings leave open the question of whether the cause of this unintuitive phenomenon is intrinsic to the model architecture and nature of single-cell data, or whether the evaluation strategy plays a role. One hypothesis is that finetuning-masking, compensation for weaker foundation representations through narrow, task-specific adaptation during partially unfrozen finetuning, might obscure the link between foundation-representation quality and pretraining scale.

### Frozen-representation evaluation reveals the pretraining scaling law obscured by standard finetuning

To test the finetuning-masking hypothesis, we repeated downstream evaluations using the same pretrained models under the fully frozen finetuning regime. We kept the input embedding layer and all encoder layers fixed and trained only the task-specific prediction head. This setting prioritises evaluating foundation representation quality, without allowing the pretrained model to adapt to task-specific data. Overall, across several classification tasks, dependence on pretraining corpus size was more pronounced than under partially unfrozen finetuning, particularly for the larger Geneformer variant.

Cell-type prediction showed the clearest contrast between the two Geneformer variants under frozen-representation evaluation (Figure 2A). Under balanced subsampling, the second-generation variant exhibited the strongest association between performance and pretraining corpus size, with mean macro-F1 increasing from 0.720 ± 0.0008 at 266k cells to 0.951 ± 0.0001 at 3.8M cells (FDR = 0.009; slope = 0.104). Performance saturation was reached at 2M cells, after which changes were modest and non-monotonic. Models pretrained on 1.1– 12M cells marginally outperformed the model pretrained on the full 34M-cell corpus (mean macro-F1 0.920 ± 0.005), whereas performance declined sharply below 1.1M cells. The first-generation variant showed a weaker response to corpus reduction, increasing mean macro-F1 from 0.783 ± 0.001 at 266k cells to 0.888 ± 0.0005 at 2M cells and 0.893 ± 0.0006 at 3.8M cells, compared with 0.882 ± 0.0007 at 34M cells (FDR = 0.010, slope = 0.057); saturation was reached at 1.1M cells. Thus, freezing exposed a substantially broader performance range in the second-generation variant, with poorer performance under severe data scarcity but stronger performance once sufficient pretraining data were available. This difference was amplified by random subsampling. For the second-generation variant, mean macro-F1 at 266k cells declined from 0.720 ± 0.0008 under balanced subsampling to 0.623 ± 0.001 under random subsampling, and remained lower at 3.8M cells (0.931 ± 0.0006 versus 0.951 ± 0.0001), although the saturation point was delayed from 2M cells to 3.8M cells. The divergence was greatest below 1.1M cells, indicating that preservation of cell-type composition was especially important for the larger second-generation variant when data were scarce. By comparison, the first-generation variant showed broadly similar performance and the same saturation point under both subsampling strategies.

Disease-status prediction showed a related, although more task-dependent, generational contrast. For heart-disease prediction, mean macro-F1 for the second-generation variant improved from 0.401 ± 0.06 at 266k cells to 0.695 ± 0.07 at 3.8M cells, compared with 0.671 ± 0.06 at 34M cells (FDR = 0.0037; slope = 0.15), with saturation reached at 3.8M cells. Models pretrained on 3.8–12M cells again slightly outperformed the full-corpus model, while performance declined sharply at the smaller corpus sizes. The first-generation variant improved more gradually, from 0.533 ± 0.06 at 266k cells to 0.634 ± 0.04 at 3.8M cells and 0.652 ± 0.05 at 34M cells (FDR = 0.0036), reaching saturation only at 7.2M cells despite the absence of a clear plateau. Random subsampling preserved the overall pattern but reduced performance, particularly for the second-generation variant: at 3.8M cells, mean macro-F1 was 0.663 ± 0.07 under random subsampling compared with 0.695 ± 0.07 under balanced subsampling, although the saturation point was unchanged. For the first-generation variant, random subsampling delayed saturation from 7.2M to 12M cells. COVID-19 prediction differed in that the second-generation variant showed a clearer scale-dependent improvement than the first-generation variant under balanced subsampling, increasing mean macro-F1 from 0.585 ± 0.3 at 266k cells to 0.818 ± 0.2 at 3.8M cells and 0.859 ± 0.1 at 34M cells (FDR = 0.019; slope = 0.092), with saturation at 3.8M cells and maximum performance at the full corpus. The first-generation variant changed only modestly, from 0.775 ± 0.1 at 266k cells to 0.809 ± 0.1 at 3.8M cells and 0.820 ± 0.1 at 34M cells (FDR = 0.051), reaching saturation at 1.1M cells. Under random subsampling, the second-generation trajectory remained broadly similar, but performance was reduced at selected intermediate scales, including 0.781 ± 0.2 at 3.8M cells compared with 0.818 ± 0.2 under balanced subsampling, and saturation was delayed until the 12M-cell corpus. For the first-generation variant, random subsampling shifted saturation in the opposite direction, from 1.1M cells to 266k cells.

Drug dosage-sensitivity prediction showed the weakest dependence on pretraining corpus size, but again revealed a stronger scaling response in the second-generation variant. Under balanced subsampling, mean macro-F1 for the second-generation variant increased from 0.831 ± 0.04 at 266k cells to 0.868 ± 0.021 at 3.8M cells and 0.874 ± 0.021 at 34M cells (FDR = 0.0036; slope = 0.027), with saturation reached at 647k cells. The first-generation variant was comparatively insensitive to scale, increasing from 0.794 ± 0.01 at 266k cells to 0.841 ± 0.02 at 314k cells before declining to 0.822 ± 0.06 at 34M cells (FDR = 0.92), and saturated at 314k cells. Random subsampling had little systematic effect on absolute performance for either variant, but shifted the saturation point earlier, from 647k to 266k cells for the second-generation variant and from 314k to 266k cells for the first-generation variant. Across all four classification tasks, the second-generation variant therefore showed stronger and more consistent sensitivity to pretraining corpus size, particularly under severe data reduction, whereas the first-generation variant generally reached saturation with less data. Most recoverable classification performance was acquired between 266k and 3.8M pretraining cells, beyond which gains were small or non-monotonic.

Perturbation-response prediction on the Norman double-perturbation benchmark showed a similar pattern, although the corresponding linear associations did not remain significant after multiple-testing correction. Under balanced subsampling, mean Pearson delta for the second-generation variant increased from 0.701 ± 0.2 at 266k cells to 0.782 ± 0.2 at 1.1M cells, where saturation was reached. The highest observed performance occurred at 12M cells (0.795 ± 0.1), exceeding performance at the full 34M-cell corpus (0.752 ± 0.2). The first-generation variant improved more gradually, from 0.630 ± 0.2 at 266k cells to 0.772 ± 0.2 at 7.2M cells, before declining to 0.678 ± 0.2 at 34M cells; its saturation threshold was reached at 2M cells. However, the associations between corpus size and mean Pearson delta were not significant after correction for either the second- or first-generation variant (FDR = 0.098 and 0.051, respectively). Random subsampling produced comparable overall trajectories to the balanced subsampling. Under the most severe data reductions, the second-generation variant retained higher mean Pearson delta after balanced subsampling, whereas random subsampling reduced performance towards that of the first-generation variant. The saturation point increased from 1.1M for the balanced subsampling to 2M for the random subsampling in the case of the second-generation variant. For the first-generation variant, however, the saturation point remained unchanged, indicating that the subsampling strategy affected absolute performance more than the corpus size required to reach saturation.

These results show that fully frozen finetuning evaluation reveals a dependence on pretraining scale that is largely obscured by the standard partially unfrozen finetuning. This dependence was strongest for the second-generation Geneformer variant and was concentrated below the pretraining corpus size of 3.8M cells, equivalent to approximately 11% of the full cellNexus corpus. Beyond this point, most downstream tasks had already reached performance saturation, and further pretraining corpus scaling produced only modest or non-monotonic changes. Balanced subsampling strategy partly mitigated the loss of foundation representation quality in the larger second-generation variant, particularly for tasks requiring fine-grained cellular discrimination. The performance saturation generally occurred later under fully frozen finetuning than under partially unfrozen finetuning, indicating that task adaptation compensates for deficiencies in foundation representations and thereby masks the underlying dependence of representation quality on pretraining scale.

### Assessment of zero-shot capability across pretraining scales

To assess how pretraining scale affects biological information encoded in the model without task-specific adaptation, we evaluated two zero-shot properties: recovery of functional gene networks (antigen-presentation gene modules) from pretrained universal gene embeddings (data-independent) ^10^, and attention-based discovery of perturbation-associated gene-regulatory relationships ^10,14^. Overall, functional gene-network recovery exhibited a clear threshold-and-saturation pattern with increasing pretraining corpus size, whereas attention-derived regulatory signals remained similar across scales but showed no consistent scaling trajectory.

### Pretraining scale affects HLA-module recovery and organisation

We first tested whether pretrained gene embeddings encoded the human leukocyte antigen (HLA) gene programme (see Methods). Under balanced subsampling, HLA gene recovery scaled with pretraining corpus size in both Geneformer variants (Figure 3A). At smaller scales of 266k to 314k cells, the second-generation variant recovered two HLA genes, and the first-generation variant recovered one. Recovery remained variable at intermediate scales: the second- and first-generation variants recovered three and four genes at 421k cells; four and two genes at 647k cells; and four and five genes at 1.1M cells, respectively. However, both variants converged on eight recovered HLA genes at 2M cells. The second-generation model reached its maximum recovery of nine genes at 3.8 million cells, whereas the first-generation model recovered eight genes at this scale before catching up with nine genes at 7.2 million cells. Recovery remained stable for both variants up to 12M cells. Notably, the 3.8M cell saturation point closely matched the saturation range observed across several fully frozen finetuning classification and perturbation-response benchmarks. At the maximum corpus size of 34M cells, recovery declined to seven genes for the second-generation variant and eight for the first-generation variant, paralleling the modest decline observed for several fully frozen finetuning downstream tasks relative to the 12M-cell models.

**Figure 3.**
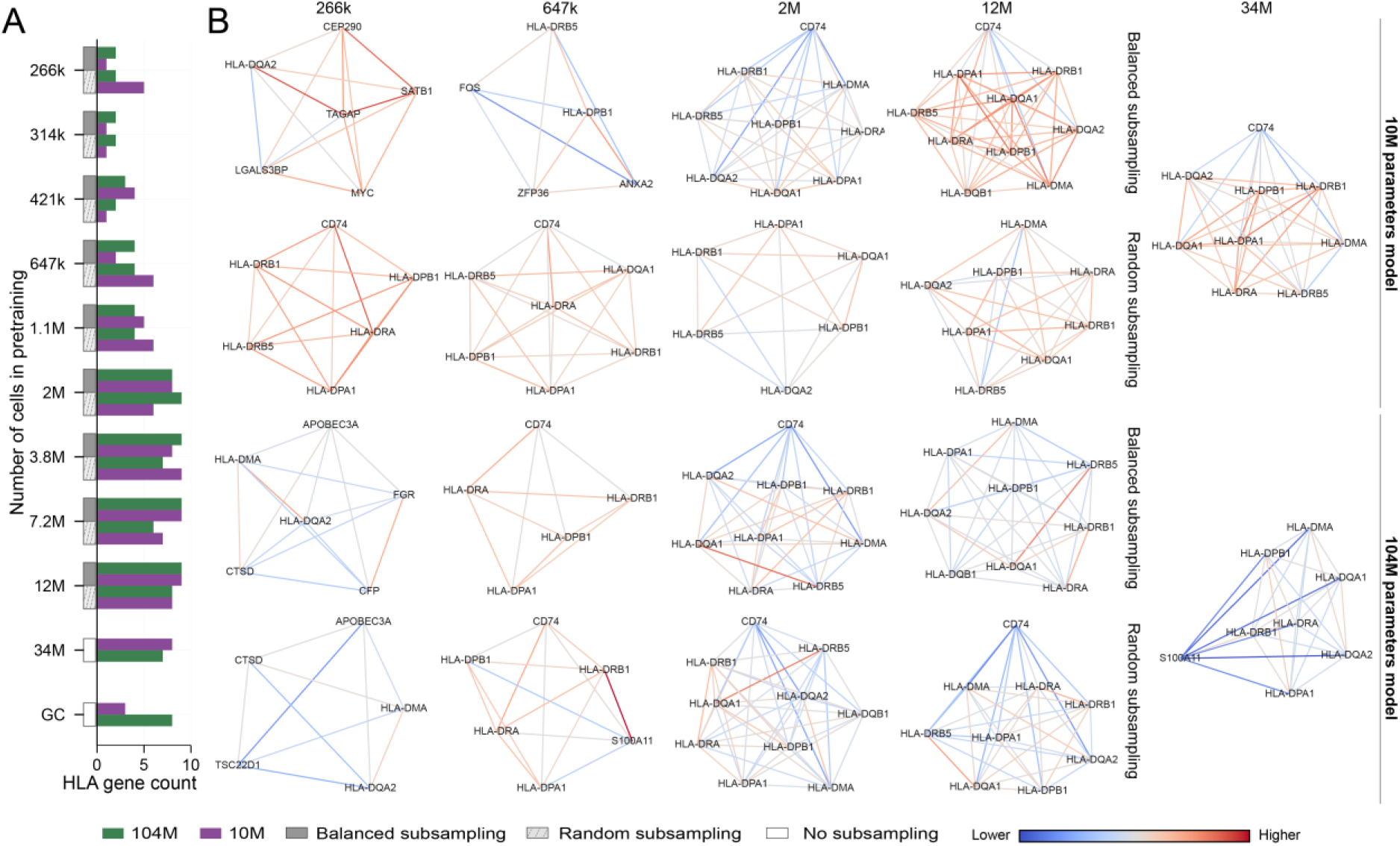
Zero-shot recovery of HLA gene modules across pretraining scales. **A**, Number of HLA genes assigned to the largest HLA-enriched Leiden cluster identified from gene-embedding similarity graphs for the first- and second-generation Geneformer variants across subsampled pretraining corpora. GC denotes the original Genecorpus trained model. **B**, Embedding-derived HLA similarity networks across corpus sizes, with edge colour indicating pairwise cosine similarity. Both model variants recovered a coherent HLA antigen-presentation module across a broad range of pretraining corpus sizes.

Compared to balanced subsampling, random subsampling produced comparable recovery at or below 1.1M cells for the second-generation variant, whereas the first-generation variant exhibited greater instability. Furthermore, between 7.2M and 12M cells, random subsampling recovered fewer HLA genes for both variants. Preserving cell-type representation therefore appeared to stabilise recovery of antigen-presentation structure before the embedding space reached its saturation. Finally, the original first-generation Geneformer pretrained on Genecorpus recovered fewer HLA genes than either the second-generation Genecorpus model or first-generation variant pretrained on comparably sized cellNexus, suggesting that both corpus composition and model size contribute to zero-shot module recovery.

We next asked whether HLA communities of similar size also showed comparable internal organisation. Pairwise cosine similarity revealed moderate-to-high cohesion across both variants and subsampling strategies (Figure 3B). The first-generation variant generally formed communities with stronger pairwise similarities than the second-generation variant, particularly at larger corpus sizes (12M and 34M cells), despite recovering similar numbers of HLA genes. HLA-gene count and within-community similarity captured complementary aspects of the embedding space: the former measured programme coverage, whereas the latter reflected the strength and heterogeneity of relationships among recovered genes.

### Attention-derived regulatory associations show no consistent scaling trend

To assess whether attention changes induced by perturbation of a transcription factor prioritised experimentally supported regulatory targets, we compared *BHLHE40*-associated attention patterns between control and perturbed cells and ranked genes by the magnitude of their perturbation-associated attention change using the Adamson CRISPR perturbation dataset ^20^. Biological support was assessed using ChIP-Atlas, a compendium of ChIP-seq-supported regulatory targets ^21^ (Figure 4A,B). Across corpus sizes, many highly ranked genes were supported by ChIP-Atlas, indicating that biologically plausible attention-derived associations were retained after substantial corpus reduction. In contrast to HLA-module recovery, attention-derived regulatory networks showed no consistent relationship with corpus size, model generation, or subsampling strategy. Under balanced subsampling, the second-generation variant identified more supported genes at 421k–647k and 3.8–7.2M cells, whereas the first-generation variant did so at 266k–314k and 1.1–2M cells. Random subsampling produced no consistent variant-specific pattern, and neither strategy showed a clear advantage. However, the second-generation variant produced smaller attention differences across both subsampling strategies than the first-generation variant, although these magnitudes cannot be compared directly because the two variants differ in attention scale and number of heads.

**Figure 4.**
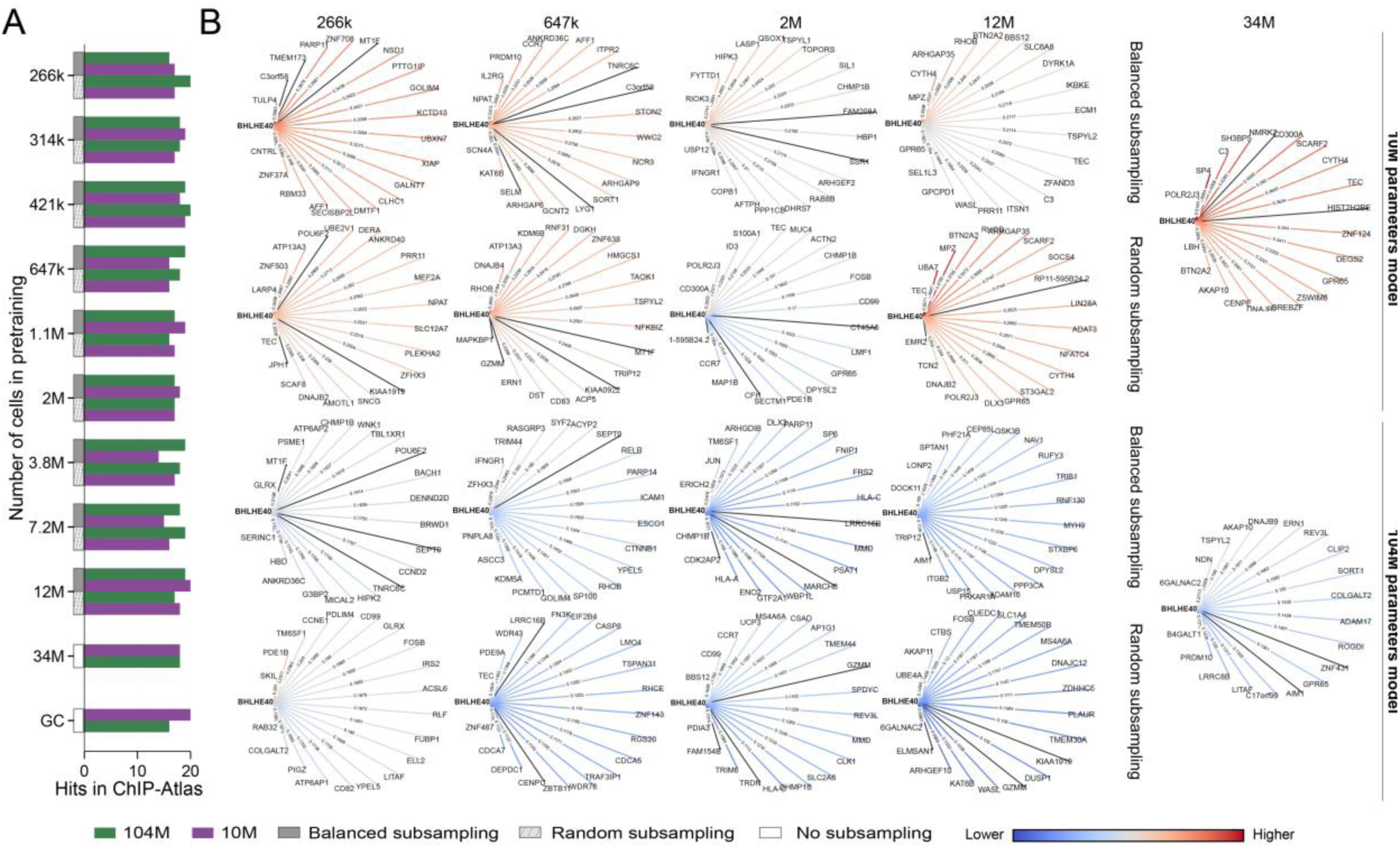
Zero-shot recovery of ChIP-Atlas-supported candidate targets from perturbation-associated attention changes. **A**, Number of ChIP-Atlas-supported targets among the top 20 genes ranked by the magnitude of attention changes following *BHLHE40* repression, inferred from last-layer attention changes. **B**, Attention-derived *BHLHE40* target networks showing the top 20 genes with the strongest repression-associated attention changes. Edges connecting *BHLHE40* to ChIP-Atlas-supported targets are coloured by attention-change magnitude, whereas edges to unsupported genes are shown in black.

These results reveal distinct effects of pretraining scale on different aspects of biological representation. HLA-module recovery followed a non-linear threshold-and-saturation pattern. Extreme corpus reduction disrupted the organisation of the antigen-presentation programme, whereas pretraining corpora of 3.8M cells or more produced stable modules with little further gain. Balanced subsampling strategy reduced instability at smaller pretraining scales. By contrast, attention-derived perturbation signals were broadly retained across pretraining corpus sizes and showed no consistent scaling trajectory. Representative results for small, intermediate, and full corpus sizes are shown in the main figure, while the complete subsampling series and the Genecorpus baseline are provided in Supplementary Figures 2 and 3.

## Discussion

Our study identifies a fundamental limitation in how cell foundation models are evaluated: conventional finetuning benchmarks can obscure improvements in biological representations acquired during pretraining. By systematically varying pretraining scale, model capacity and evaluation strategy, we demonstrate that task-specific adaptation can compensate for weaker pretrained representations, a phenomenon we term finetuning masking. This effect was particularly pronounced in the second-generation Geneformer model, where fully frozen evaluation revealed a stronger dependence on pretraining scale. These findings establish that downstream performance saturation cannot reliably determine whether further pretraining improves the underlying biological representations.

The fully frozen benchmark allowed us to disentangle representation saturation from data-starvation regimes. The stronger effect of corpus reduction in the second-generation variant highlights the data-to-model-size relationship. The larger variant showed greater sensitivity to data starvation under fully frozen finetuning evaluation, while also achieving stronger performance in data-rich settings. This broader dynamic range is consistent with a model that can exploit additional pretraining data but requires sufficient exposure to realise that capacity. The lower sensitivity of the first-generation variant should therefore not be taken to indicate uniformly stronger foundation representations. Its more limited capacity may constrain both the structure it can extract from additional data and the extent to which its performance deteriorates when data are removed. A pretraining-scale benchmark in a fully frozen setting could therefore be used to estimate the effective model size given pretraining size.

The comparison between balanced and random subsampling also indicates that corpus size alone provides an incomplete description of pretraining exposure. Preserving diversity across donors, cell types, and biological contexts became more consequential when downstream prediction depended directly on the fixed foundation representation under fully frozen finetuning. Under partially unfrozen finetuning, these differences were less apparent, consistent with task adaptation compensating for losses in biological coverage. Corpus-reduction experiments should therefore distinguish reductions in redundant cell numbers from reductions in distinct biological contexts. This distinction is especially important in single-cell atlases, where millions of cells may originate from a comparatively limited number of donors, tissues or disease settings. Benchmarking future generations of models will further clarify the extent to which learning saturation is linked to model size, architecture, or the intrinsic redundancy of single-cell molecular data.

The zero-shot analyses provided an independent perspective on how pretraining data shape biological representations. Recovery of the HLA module broadly tracked the fully frozen finetuning benchmarks, suggesting that the scale dependence detected under these benchmarks reflects changes in the organisation of the foundation embedding space and not only improved separability for individual supervised tasks. The lack of a comparable scaling trajectory for the attention-based *BHLHE40* analysis indicates that distinct biological probes interrogate different properties of the model. Gene-module recovery evaluates whether functionally related genes occupy a coherent region of the input embedding space, whereas attention-based perturbation analysis captures context-dependent associations after token contextualisation. Some regulatory-like signals may therefore emerge from relatively small corpora or may be less sensitive to the global organisation of the embedding space.

Several considerations define the scope of this study. Our analyses focused on two Geneformer variants, and evaluation across additional single-cell foundation-model families will be required to establish the generality of the observed scaling behaviour. The tested parameter and corpus ranges also do not define the upper limits at which further increases in model size or pretraining corpus may remain beneficial. In addition, our saturation criterion should be interpreted as an empirical threshold within the tested range, not as a universal data requirement.

These findings have implications for how single-cell foundation models are evaluated, compared and developed. Conventional standard finetuning benchmarks primarily measure whether a model can be adapted to solve a predefined task, while providing limited information about the quality of the representation available before adaptation. Fully frozen finetuning evaluation offers a stricter test of transferable information, and zero-shot analyses assess whether biologically meaningful structure can be accessed without task-specific encoder updates. We therefore propose that foundation-model evaluation combine standard finetuning, fully frozen finetuning, data-starvation curves and zero-shot biological inference. Reporting how performance changes across these protocols would help distinguish models that adapt effectively to narrow supervised objectives from those that encode broadly transferable biological structure. Ultimately, progress in cell foundation models will require evaluation strategies that measure the biological information acquired during pretraining and its transferability across increasingly complex and open-ended biological problems.

## Methods

### Pretraining data and subsampling strategies

To evaluate the effect of pretraining corpus size, we used cellNexus ^17^ as the source data. cellNexus provides standard quality-control filtering, including removal of contaminated empty droplets, low-quality cells and predicted doublets. Our curated collection contains over 34 million cells from 172 tissues, spanning healthy donors and diverse disease contexts, and therefore provided broad coverage of the human single-cell transcriptomic landscape.

### Balanced subsampling

Reducing a large single-cell pretraining corpus by naive subsampling can distort both biological and study-level structure by eliminating rare samples, donors, datasets or cell populations and over-representing abundant cell types. To limit these effects, we developed a balanced subsampling strategy that retained representation from each metadata-defined group while approximately preserving the cell-type composition within that group (Figure 1A, B; Supplementary Figure 1A,B). This design was intended to preserve biological diversity as corpus size decreased. In particular, it reduced the likelihood that the smallest corpora would be dominated by a limited number of large samples or common cell-type populations. This distinction is important because prior work suggests that diversity can be more informative for foundation model pretraining than increasing the volume of biologically redundant data ^15^. By retaining sample-level coverage while reducing the number of cells contributed by each sample, the procedure also helped separate the effect of corpus size from changes in cohort composition. Thus, corpus reduction preferentially removed redundant observations before eliminating distinct biological contexts.

The full corpus was partitioned into *G* disjoint groups defined by the metadata tuple:

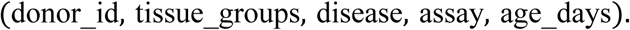

Each group therefore represented a distinct combination of biological and technical attributes, and subsampling was performed independently within each group.

For a requested corpus size *n*, near-equal target quotas were assigned across groups to define a group-specific sampling plan. These quotas were treated as approximate targets rather than strict upper bounds. Within each group, cells were sampled without replacement, while ensuring that every cell type represented in the cell_type_unified_ensemble metadata field contributed at least one cell. Because preserving cell-type coverage was prioritised over exact adherence to the assigned quotas, the number of cells retained from a given group could deviate from its nominal allocation. Groups containing many distinct cell types could therefore contribute more cells than their assigned quota, whereas groups with fewer available cells could contribute less. As a result, the final subsampled corpus sizes approximated the requested targets while preserving broad group-level representation and cell-type diversity. All sampling was performed using a fixed random seed to ensure reproducibility.

### Random subsampling

To provide a comparator that did not explicitly preserve biological or study structure, we also generated randomly subsampled corpora. Cells were sampled uniformly without replacement from the full cellNexus corpus, without stratification by any metadata (Figure 1A, B). Randomly subsampled datasets were constructed to match the corresponding balanced-subsampling corpus sizes, enabling direct comparison of the effects of corpus size and corpus composition.

### Pretraining of foundation models

To assess how pretraining corpus size interacts with model capacity, we pretrained the second-generation (104-million parameters with 12 encoder layers and 12 attention heads) ^15^ and first-generation (10-million parameters with 6 encoder layers and 4 attention heads) ^8^ Geneformer. These models provide two substantially different capacities within a shared architectural framework, enabling controlled comparison under matched pretraining and downstream evaluation protocols. This design allowed us to test whether the larger architecture learned stronger foundation representations and whether any advantage depended on the amount of pretraining data available. The pretraining design is summarised in Figure 1A and the reduction in computational cost across progressively subsampled corpora is shown in Figure 1C.

Geneformer represents each cell as a rank-ordered sequence of expressed genes during tokenisation. In this study, tokenisation followed the procedures described for the respective Geneformer implementations ^8,15^. Both variants were pretrained for 3 epochs. The principal architectural and optimisation hyperparameters were as follows. For the second-generation variant: non-linear activation function, ReLU; dropout probability for fully connected layers, 0.1; dropout ratio for attention probabilities, 0.1; standard deviation of the initialiser for weight matrices, 0.02; epsilon for layer-normalisation layers, 1 × 10^−1^^2^; feed-forward size, 3,072; embedding dimension, 768; weight decay, 0.044; learning-rate scheduler, cosine; learning-rate, 0.0002; warmup steps, 10,000; batch size, 18. For the first-generation variant: non-linear activation function, ReLU; dropout probability for fully connected layers, 0.02; dropout ratio for attention probabilities, 0.02; standard deviation of the initialiser for weight matrices, 0.02; epsilon for layer-normalisation layers, 1 × 10^−1^^2^; feed-forward size, 512; embedding dimension, 256; weight decay, 0.001; learning-rate scheduler, linear; learning-rate, 0.001; warmup steps, 10,000; batch size, 12. The second-generation models were pretrained on a compute node equipped with four NVIDIA H200 GPUs with 141GB memory each, whereas the first-generation models were pretrained on NVIDIA A100 GPUs with 40GB memory.

### Partially unfrozen and fully frozen finetuning regimes

For each downstream task and Geneformer variant, we evaluated two adaptation regimes: partially unfrozen finetuning and fully frozen finetuning. In partially unfrozen finetuning, the task-specific prediction head, input embedding layer and upper encoder layers were updated during finetuning. Specifically, the top two encoder layers were unfrozen for the second-generation model, whereas the top three layers were unfrozen for the first-generation model. The number of trainable layers was selected empirically following the procedure described previously ^8^. This setting reflects standard task adaptation, in which both the prediction head and high-level foundation representations can adjust to the downstream objective. In fully frozen finetuning, the input embedding layer and all encoder layers were held fixed, and only the task-specific prediction head was trained. This setting isolates the information already encoded during pretraining by preventing task-specific adaptation of the foundation representation.

More formally, let the pretrained model comprise an input embedding layer, *L* stacked encoder layers and a task-specific prediction head, with pretrained parameters denoted collectively by Θ. In the partially unfrozen finetuning, the prediction head and the top *k* encoder layers were trained along with the embedding layer:

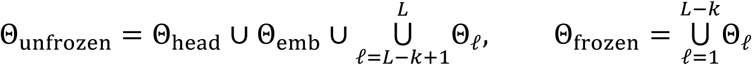

where *k* = 2 for the second-generation variant and *k* = 3 for the first-generation variant. In empirical tests, unfreezing more than two encoder layers during finetuning of the second-generation variant led to overfitting. We therefore used different values of *k* for both variants. In the fully frozen finetuning, only the task-specific prediction head parameters were updated:

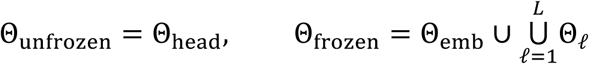

These two finetuning regimes capture distinct ways in which pretraining corpus scale can influence downstream performance. In the partially unfrozen setting, performance reflects how effectively higher-level foundation representations can be adapted to a new task. By contrast, fully frozen finetuning performance primarily reflects the accessibility of task-relevant information already encoded in the foundation representation. Comparing these finetuning regimes therefore allowed us to distinguish apparent downstream performance saturation driven by task adaptation from saturation of the underlying foundation representation.

### Finetuning on downstream tasks

#### Classification tasks

All classification tasks were evaluated using a standardised finetuning framework adapted from the original Geneformer studies ^8,15^. To enable controlled comparisons across tasks, pretraining corpus sizes and subsampling strategies, finetuning hyperparameters were fixed within each Geneformer variant rather than optimised separately for individual corpus scales. Hyperparameter settings were selected empirically.

The second-generation variant was finetuned for two epochs using a learning rate of 3.55 × 10^−4^, cosine learning-rate scheduling, 500 warm-up steps, weight decay of 0.05 and a per-device batch size of 8. The first-generation variant was finetuned for ten epochs using a learning rate of 8.04 × 10^−4^, polynomial learning-rate scheduling, 1,812 warm-up steps, weight decay of 0.258828 and a per-device batch size of 64. For both variants, the random seed was fixed to 73. Different numbers finetuning epochs were used because empirical testing showed that training the second-generation variant for more than two epochs resulted in overfitting. Validation loss was used for checkpointing after each epoch and the checkpoint with the lowest validation loss was retained for final evaluation.

Performance was evaluated using five-fold cross-validation. For each model, task and pretraining corpus, macro-F1 was calculated independently for each fold, consistent with the literature ^8,10,14,16^, and summarised as the mean and standard deviation across folds. More formally, F1 for a particular class *c* is defined as:

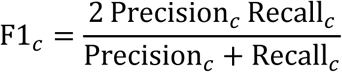

where

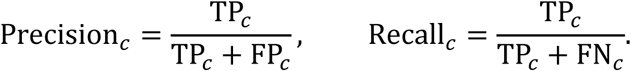

Here, TP denotes the number of true positives, FP represents the number of false positives, and FN is the number of false negatives. Macro-F1 for a particular fold is then defined as:

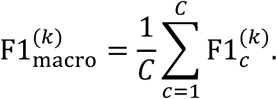

Mean macro-F1 is an arithmetic average across all five folds:

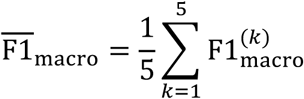

Fold construction was task-specific and designed to prevent leakage of biological or study-level information, as described below. For the performance saturation threshold, we identify the smallest corpus size achieving 95% of the maximum observed performance (after spline smoothing) for a particular model variant and subsampling strategy.

To evaluate the model’s ability for cell-type discrimination, we constructed a subset of cellNexus comprising 593,438 cells from 51 cell types. Cells were assigned uniformly at random to five folds. For each fold, the model was trained on four folds (training partitions) and evaluated on the held-out fold (validation partition).

The heart-disease prediction dataset was obtained from Chaffin et al. ^22^ and is tokenised following the original Geneformer studies ^8,15^. Five-fold cross-validation was performed at the patient level, such that patients assigned to the training and validation sets were mutually exclusive within each fold. This design prevented leakage of patient-specific transcriptional profiles across splits.

The COVID-19 benchmark was constructed from a cellNexus subset comprising 20 independent datasets, including 159,738 normal and 158,956 COVID-19 cells. Cross-validation was performed at the dataset level: within each fold, all cells from a given dataset were assigned exclusively to either the training or validation partition. This design prevented dataset-specific technical and cohort signals from being shared across partitions and provided a stringent evaluation of cross-dataset generalisation.

We evaluated the ability of the models to distinguish dosage-sensitive from dosage-insensitive transcription factors ^8^. Labels were derived from previously curated dosage-sensitivity annotations ^23–25^. Models were finetuned using the 50,000-cell dataset released with the original Geneformer study ^8^. Five-fold cross-validation was performed by partitioning transcription factors, such that genes assigned to the training and validation partitions were mutually exclusive within each fold.

#### Perturbation-response prediction task

We evaluated both Geneformer variants on the double-gene perturbation response prediction using the Norman dataset ^18^, accessed in its preprocessed form through the GEARS package ^26^. We adapted the Geneformer benchmarking workflow from Ahlmann-Eltze et al. ^19^, available on GitHub (linear_perturbation_prediction_Paper). Whereas Ahlmann-Eltze et al. averaged performance across five random perturbation-level train–test splits, we used a single fixed split shared across all model configurations. Consistent with prior work ^19^, perturbation prediction performance was evaluated on the 1,000 most highly expressed genes in the control condition. We use Pearson delta as a performance metric following Cui et al. ^10^ and Ahlmann-Eltze et al. ^19^, which is defined as

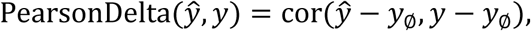

where *y*_∅_ denotes the mean expression profile of the control condition. For completeness, we also calculated the *L*_2_ distance metric following Ahlmann-Eltze et al. ^19^ as:

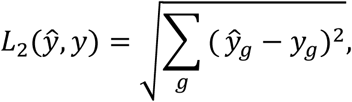

where ŷ and *y* denote the predicted and observed expressions, respectively, and the sum is over the selected 1,000 genes. However, in the main text, we only focus on Pearson delta and report the *L*_2_ distance in the Supplementary Table 2. For each experimental configuration, we report the mean and standard deviation of the per-perturbation scores across the held-out test split.

#### Statistical testing of corpus-size effects

To determine whether downstream performance varied systematically with pretraining corpus size, we fitted ordinary least-squares regression models separately for each model size, finetuning regime, sampling strategy, downstream task, and evaluation metric:

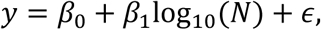

where *y* is the mean downstream performance for a given pretraining corpus size and *N* is the number of cells in the pretraining corpus. The slope *β*_1_, therefore, represents the expected absolute change in performance associated with a tenfold increase in pretraining corpus size.

We tested the null hypothesis *β*_1_ = 0 for each fitted line and adjusted the resulting *P* values across all tested associations using the Benjamini-Hochberg false discovery rate (FDR) procedure. To determine whether an observed association was not only statistically supported but also exceeded a predefined minimum effect size, we additionally performed the one-sided minimum-effect test

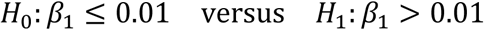

The threshold corresponds to an absolute performance increase of 0.01 per tenfold increase in pretraining corpus size, equivalent to one percentage point for metrics expressed on a 0–1 scale. The *P* values from the minimum-effect tests were adjusted separately across the same set of associations using the Benjamini–Hochberg procedure. Reported FDR values therefore quantify either evidence of any association with corpus size or, where explicitly stated, evidence that the positive association exceeded the predefined minimum-effect threshold (Supplementary Table 1).

### Zero-shot analysis

#### Gene embeddings-based community detection

For zero-shot analysis of gene-level structure, we extracted the universal, non-contextual gene embeddings from the input embedding layer of each pretrained Geneformer model. Because these embeddings are learned during pretraining and do not depend on the downstream dataset, they provide a direct representation of gene relationships encoded by the model. We constructed a *k*-nearest-neighbour graph over genes using cosine similarity between embedding vectors and applied Leiden community detection at a resolution of 20, following the general approach of Cui et al. ^10^. Leiden clusters were interpreted as candidate gene programmes, and clusters containing at least five genes were retained for the analysis. To quantify recovery of the human leukocyte antigen (HLA) programme, we identified the retained Leiden clusters containing the largest number of HLA-related genes. If no retained cluster contained an HLA gene, recovery was recorded as zero. HLA-module recovery was quantified as the number of HLA genes assigned to this community. To assess HLA-cluster cohesion, we calculated pairwise cosine similarity among all genes in the selected HLA-enriched cluster. The resulting weighted gene–gene network was visualised with genes as nodes and cosine similarities as edge weights.

#### Attention-derived gene regulatory network

For attention-based target-gene analysis of perturbation-associated gene interactions, we followed the general framework of Cui et al. ^10^. We extracted attention matrices from the final encoder layer for control cells and cells perturbed for BHLHE40. Attention scores were rank-normalised sequentially across rows and columns, averaged across attention heads and then aggregated across cells to generate gene–gene interaction maps, separate for control and perturbed cells. Perturbation-associated changes were quantified from the difference between perturbed and control attention maps. Genes were ranked using values in the column corresponding to *BHLHE40*, and the top 20 genes were retained as the most strongly affected candidates. These genes were compared against experimentally validated *BHLHE40* targets from ChIP-Atlas. Overlapping genes were classified as hits, quantifying how effectively attention-derived rankings recovered independently supported regulatory relationships.

The study contains supplementary material available online.

We thank L. Coin and F. Li for their valuable feedback. We acknowledge the computational resources and services provided by the National Computational Infrastructure (NCI), supported by the Australian Government, through the Australian BioCommons Leadership Share (ABLeS), which enabled this study. We also acknowledge the computational support provided by the Adelaide University High-Performance Computing service.

## Declarations

### Consent for publication

All authors have approved the manuscript and consent to its publication.

### Data availability

The data used in this study are publicly available through cellNexus at https://cellnexus.org. Owing to their large size, the tokenised pretraining datasets are available from the corresponding authors upon request. The pretrained models and tokenised task-specific datasets are available at Google Drive.

### Materials availability

Not applicable.

### Code availability

The code used in this paper is available at https://github.com/MangiolaLaboratory/shakeel_et_al_FM_pretraining.

### Author contributions

M.H.S. and S.M. developed the concept and contributed to the conception and design of the study. M.H.S. implemented the experiments. M.H.S. and S.M. analysed the computational experiments and drafted the manuscript. M.S. contributed to dataset curation.

