## Supplementary Figures for "Re-evaluating the benefit of training scale in cell foundation models"

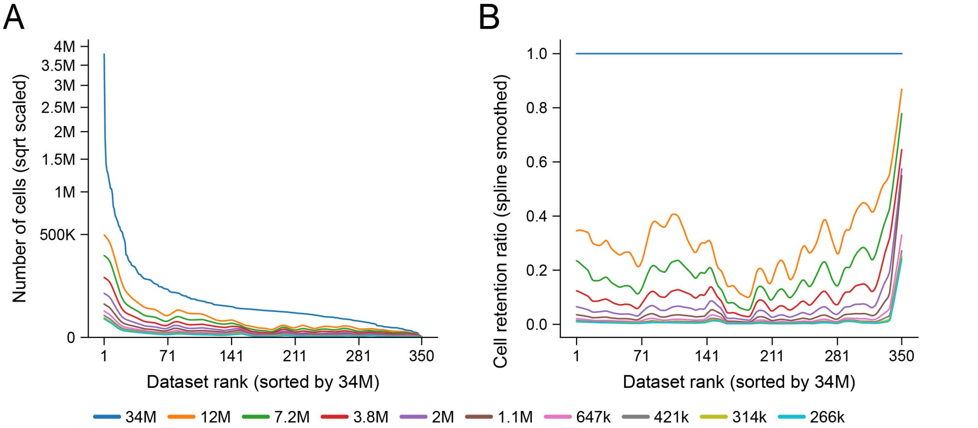


**Supplementary Figure S1:** Balanced subsampling and computational scaling. **A:** Distribution of dataset sizes across subsampled levels of cellNexus, plotted on a square-root scale. Larger datasets were reduced more strongly, while smaller datasets were comparitively preserved. **B:** Cell-retention ratio of each dataset relative to the full 34M-cell corpus for each subsampling level, showing preferential retention of cells from smaller datasets.

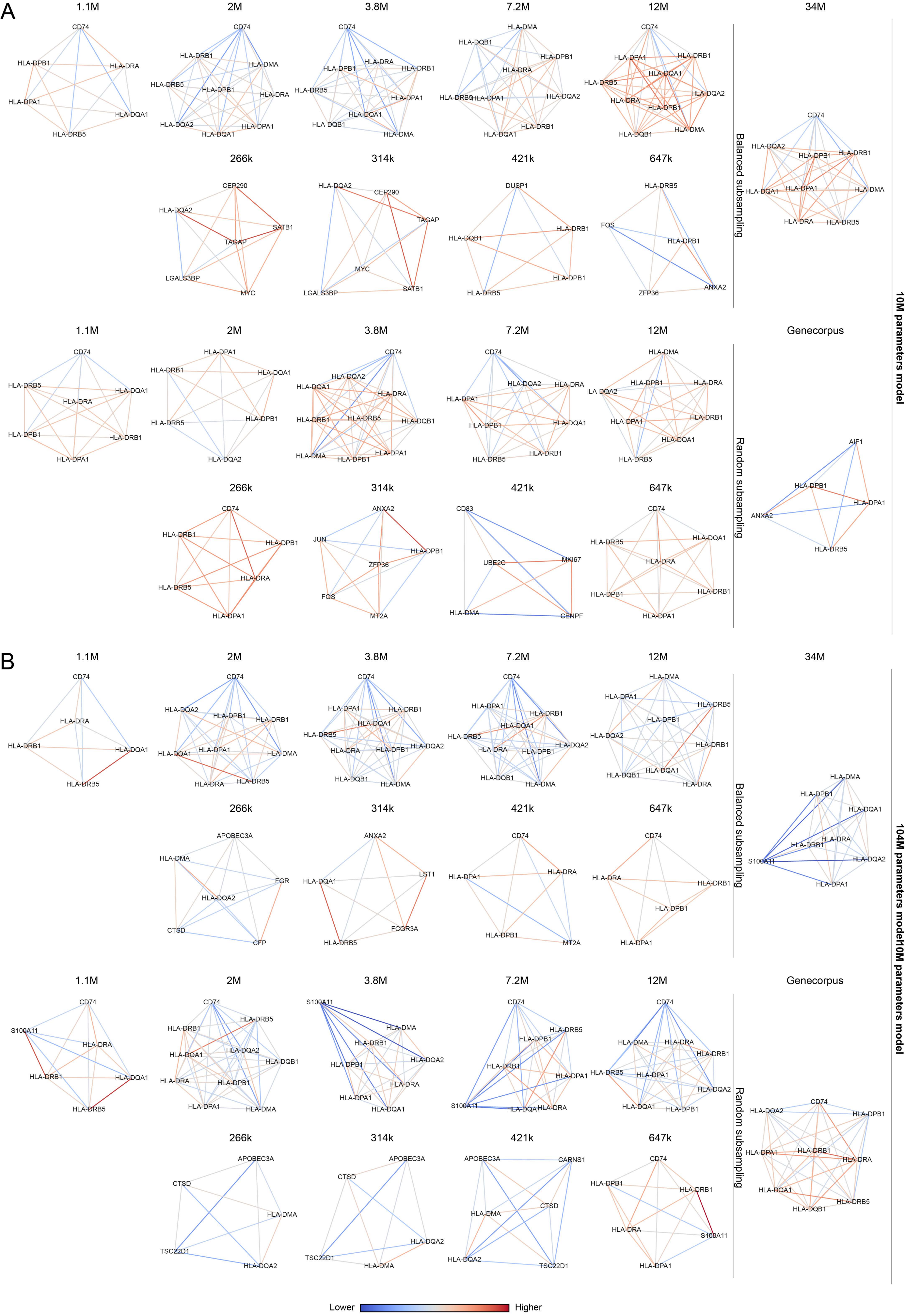


**Supplementary Figure S2:** HLA-genes communities constructed with Leiden clustering. **A:** Largest gene cluster containing HLA genes identified from gene-embeddings similarity networks for the first-generation Geneformer across all subsampling variants and Genecorpus-pretrained model. **B:** Largest gene cluster containing HLA genes identified from gene-embedding similarity networks for the second-generation Geneformer models across all subsampling variants and the Genecorpus-pretrained model.

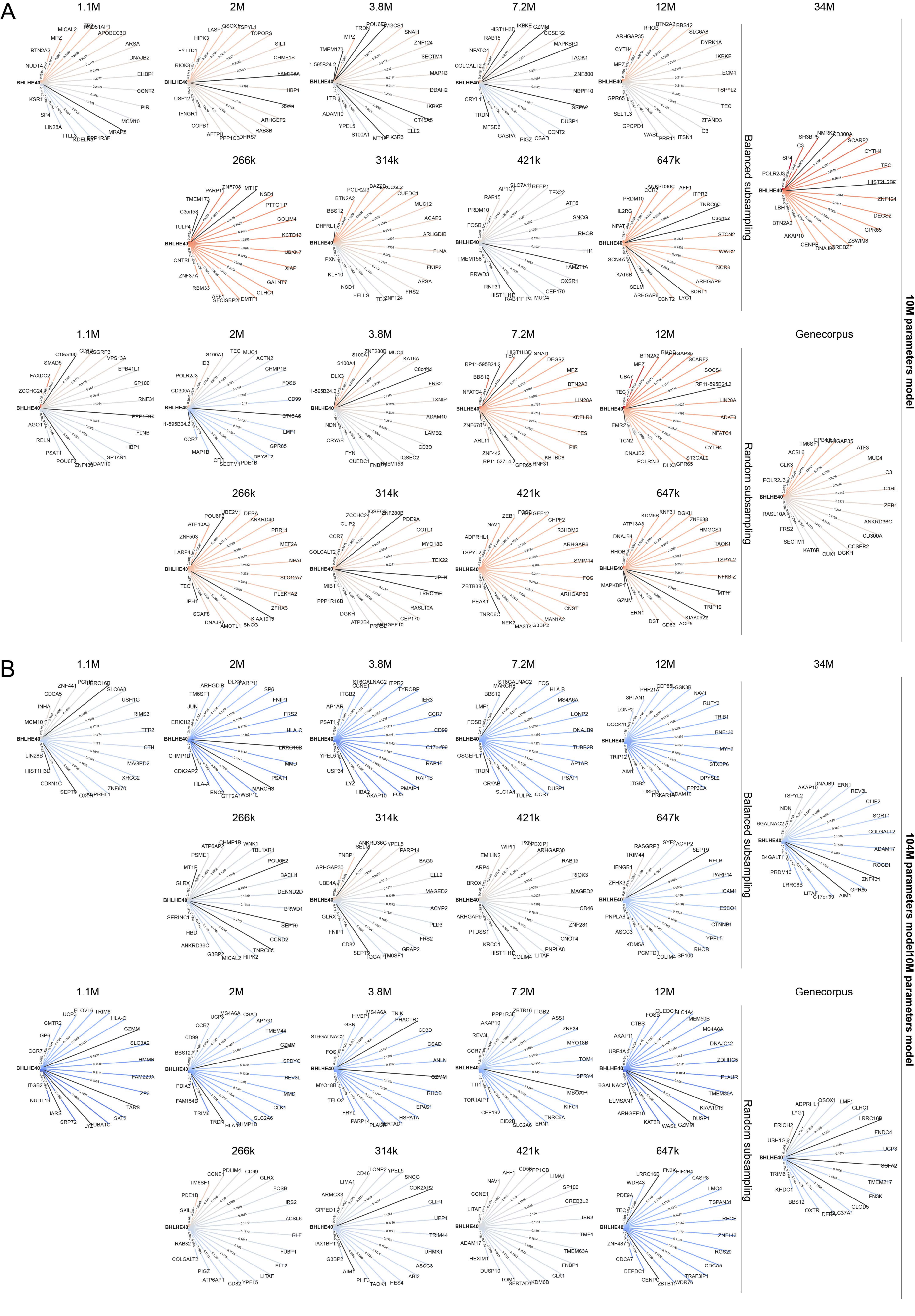


**Supplementary Figure S3:** Attention-derived gene networks and ChIP-Atlas-supported BHLHE40 targets. **A:** Attention-derived gene networks for the first-generation Geneformer models across all subsampling variants and the Genecorpus-pretrained model. **B:** Attention-derived gene networks for the second-generation Geneformer models across all subsampling variants and the Genecorpus-pretrained model.
