## Supplementary Table 2 for "Re-evaluating the benefit of training scale in cell foundation models"

| Model Variant | Freezing Strategy | Task | Subsampling Strategy | Pretraining Cells | L2 Distance | Pearson Delta | L2 Distance STD | Pearson Delta STD | n_perturbations |
|---|---|---|---|---|---|---|---|---|---|
| 10M | Fully Frozen | Norman Perturbation | balanced | 266k | 4.672996114160089 | 0.6308701363397691 | 1.921306585525947 | 0.2590156267990274 | 62 |
| 10M | Fully Frozen | Norman Perturbation | balanced | 314k | 4.7973606878813495 | 0.5933867742022598 | 1.9217369042867092 | 0.24023793004394417 | 62 |
| 10M | Fully Frozen | Norman Perturbation | balanced | 421k | 4.648299949668511 | 0.63085542 | 1.8950482178413255 | 0.2548811642645725 | 62 |
| 10M | Fully Frozen | Norman Perturbation | balanced | 647k | 4.649846524244384 | 0.6280769307905532 | 1.906456798917607 | 0.2394899773266156 | 62 |
| 10M | Fully Frozen | Norman Perturbation | balanced | 1.1M | 4.328883933602717 | 0.687975784 | 1.7612908886324463 | 0.24647211572595973 | 62 |
| 10M | Fully Frozen | Norman Perturbation | balanced | 2M | 4.165668824221984 | 0.7072130076484917 | 1.5303173318596714 | 0.248926195 | 62 |
| 10M | Fully Frozen | Norman Perturbation | balanced | 3.8M | 4.283306284978173 | 0.6983636797095132 | 1.753279073058677 | 0.23847667200322625 | 62 |
| 10M | Fully Frozen | Norman Perturbation | balanced | 7.2M | 3.838235776346928 | 0.772802969 | 1.4990345042574869 | 0.20366910877122463 | 62 |
| 10M | Fully Frozen | Norman Perturbation | balanced | 12M | 4.2928193776642765 | 0.6893596373241421 | 1.743500473666706 | 0.230629291818452 4 | 62 |
| 10M | Fully Frozen | Norman Perturbation | full corpus | 34M | 4.430010824858426 | 0.6786497750351247 | 1.7534722282222155 | 0.22081731201431862 | 62 |
| 10M | Fully Frozen | Norman Perturbation | random | 266k | 4.447517612787685 | 0.6684334602685893 | 1.7080303602005518 | 0.26018161353434877 | 62 |
| 10M | Fully Frozen | Norman Perturbation | random | 314k | 4.3664513622197765 | 0.6721947738830754 | 1.808831214173191 | 0.2653972017705428 | 62 |
| 10M | Fully Frozen | Norman Perturbation | random | 421k | 4.6441251405574935 | 0.6196099927250136 | 1.8303095416276234 | 0.2422861775766881 | 62 |
| 10M | Fully Frozen | Norman Perturbation | random | 647k | 4.272122842048296 | 0.6945665964151138 | 1.6958142067626139 | 0.2510656354871734 | 62 |
| 10M | Fully Frozen | Norman Perturbation | random | 1.1M | 4.277749503 | 0.6974208304428808 | 1.7395490258885276 | 0.2609739886082542 | 62 |
| 10M | Fully Frozen | Norman Perturbation | random | 2M | 4.237646936373616 | 0.7071650017666563 | 1.7970350956833476 | 0.24159843945470683 | 62 |
| 10M | Fully Frozen | Norman Perturbation | random | 3.8M | 3.8785244303800512 | 0.7468329684134103 | 1.7921348134205652 | 0.24283469614222034 | 62 |
| 10M | Fully Frozen | Norman Perturbation | random | 7.2M | 3.8218448514501797 | 0.7690411392363292 | 1.621065727907379 | 0.21380961346105326 | 62 |
| 10M | Fully Frozen | Norman Perturbation | random | 12M | 4.2714075582295425 | 0.7093394542030829 | 1.847178172644742 | 0.2374705733851106 | 62 |
| 10M | Partially Unfrozen | Norman Perturbation | balanced | 266k | 4.318859267476313 | 0.6790134456716385 | 1.6666765054629848 | 0.25012932750409095 | 62 |
| 10M | Partially Unfrozen | Norman Perturbation | balanced | 314k | 4.336823300810157 | 0.6729073879831254 | 1.617295181399122 | 0.23599904592906767 | 62 |
| 10M | Partially Unfrozen | Norman Perturbation | balanced | 421k | 4.375878613990833 | 0.6582060039343182 | 1.657567006504564 | 0.26482815658418457 | 62 |
| 10M | Partially Unfrozen | Norman Perturbation | balanced | 647k | 4.111626174606266 | 0.7283210032119057 | 1.689642359 | 0.21390730432500157 | 62 |
| 10M | Partially Unfrozen | Norman Perturbation | balanced | 1.1M | 3.688697696 | 0.7605778355371239 | 1.332624904824807 | 0.22017320158733528 | 62 |
| 10M | Partially Unfrozen | Norman Perturbation | balanced | 2M | 3.4643542492686525 | 0.7878905800616742 | 1.241199538182511 | 0.22318643957566006 | 62 |
| 10M | Partially Unfrozen | Norman Perturbation | balanced | 3.8M | 3.5162556074613254 | 0.7960871630912338 | 1.3868874726588867 | 0.2092295872050262 | 62 |
| 10M | Partially Unfrozen | Norman Perturbation | balanced | 7.2M | 3.408708482 | 0.7962043360305944 | 1.272043369096966 | 0.19384041708029198 | 62 |
| 10M | Partially Unfrozen | Norman Perturbation | balanced | 12M | 3.612548406728803 | 0.7947913367546287 | 1.375650809918034 | 0.17529041120692584 | 62 |
| 10M | Partially Unfrozen | Norman Perturbation | full corpus | 34M | 3.502084078358066 | 0.8000337138619382 | 1.3768874913794533 | 0.16818051717755683 | 62 |
| 10M | Partially Unfrozen | Norman Perturbation | random | 266k | 4.244804355563763 | 0.692380839 | 1.5576783047294658 | 0.23970846392757927 | 62 |
| 10M | Partially Unfrozen | Norman Perturbation | random | 314k | 4.130905648234845 | 0.6963228076370065 | 1.491407076773341 | 0.21281230929139122 | 62 |
| 10M | Partially Unfrozen | Norman Perturbation | random | 421k | 4.215457705705048 | 0.6956487855549848 | 1.4871191954718563 | 0.20313719081059675 | 62 |
| 10M | Partially Unfrozen | Norman Perturbation | random | 647k | 3.9411702583389876 | 0.7354163373481212 | 1.4512021897287544 | 0.2329646203726167 | 62 |
| 10M | Partially Unfrozen | Norman Perturbation | random | 1.1M | 3.451322017937622 | 0.7787730680957691 | 1.1122341327629333 | 0.2420361017995623 | 62 |
| 10M | Partially Unfrozen | Norman Perturbation | random | 2M | 3.426658420364479 | 0.8010668898962701 | 1.2865599034305042 | 0.19823404700175634 | 62 |
| 10M | Partially Unfrozen | Norman Perturbation | random | 3.8M | 3.5120643667659546 | 0.7898301038391417 | 1.334169447 | 0.21816607503840044 | 62 |
| 10M | Partially Unfrozen | Norman Perturbation | random | 7.2M | 3.3866377477958243 | 0.8142704650962735 | 1.3941267724694513 | 0.15176060037265543 | 62 |
| 10M | Partially Unfrozen | Norman Perturbation | random | 12M | 3.374594561833723 | 0.8100016959167045 | 1.4391708842558601 | 0.20446323217988907 | 62 |
| 104M | Fully Frozen | Norman Perturbation | balanced | 266k | 4.220709353878193 | 0.7012933661447701 | 1.7188806557607017 | 0.23696057937187548 | 62 |

| | | | | | | | | | |
|---|---|---|---|---|---|---|---|---|---|
| 104M | Fully Frozen | Norman Perturbation | balanced | 314k | 4.315285454 | 0.6805278292049115 | 1.7133368305581107 | 0.2369886843677212 | 62 |
| 104M | Fully Frozen | Norman Perturbation | balanced | 421k | 3.808919189 | 0.7472717757863776 | 1.385833734776454 | 0.21152950049492522 | 62 |
| 104M | Fully Frozen | Norman Perturbation | balanced | 647k | 4.289645147497602 | 0.6959856845403692 | 1.695729522066782 | 0.23121154748328054 | 62 |
| 104M | Fully Frozen | Norman Perturbation | balanced | 1.1M | 3.6653948313286797 | 0.7821527726760661 | 1.5192884847878303 | 0.21313607808182755 | 62 |
| 104M | Fully Frozen | Norman Perturbation | balanced | 2M | 3.6473014060948303 | 0.7880186750767385 | 1.3935270834873037 | 0.19970876093448353 | 62 |
| 104M | Fully Frozen | Norman Perturbation | balanced | 3.8M | 3.8367671062798343 | 0.7552398541493601 | 1.4781651647200171 | 0.2131670782448365 | 62 |
| 104M | Fully Frozen | Norman Perturbation | balanced | 7.2M | 3.9025322928779422 | 0.74997154 | 1.421835585021093 | 0.21259140688752867 | 62 |
| 104M | Fully Frozen | Norman Perturbation | balanced | 12M | 3.585017992470993 | 0.7956527004475072 | 1.3138020166199382 | 0.17016254403428505 | 62 |
| 104M | Fully Frozen | Norman Perturbation | full corpus | 34M | 3.858360908075879 | 0.7523425123107655 | 1.458411253600029 | 0.21285192361637115 | 62 |
| 104M | Fully Frozen | Norman Perturbation | random | 266k | 4.339501341337409 | 0.6709866839171714 | 1.75016013 | 0.25820623368698314 | 62 |
| 104M | Fully Frozen | Norman Perturbation | random | 314k | 4.500567007029992 | 0.6597532794717124 | 1.8314886908240193 | 0.25175591837796596 | 62 |
| 104M | Fully Frozen | Norman Perturbation | random | 421k | 4.445347484 | 0.6550578478740945 | 1.6990182623678756 | 0.24248499331401702 | 62 |
| 104M | Fully Frozen | Norman Perturbation | random | 647k | 4.294410737003209 | 0.6959241646159957 | 1.6732110298542442 | 0.24278512447249717 | 62 |
| 104M | Fully Frozen | Norman Perturbation | random | 1.1M | 3.8454942655176465 | 0.7538324782139608 | 1.4184357174853193 | 0.22122892644897693 | 62 |
| 104M | Fully Frozen | Norman Perturbation | random | 2M | 3.9344132018816267 | 0.740765809 | 1.5189169530866498 | 0.23665570909335504 | 62 |
| 104M | Fully Frozen | Norman Perturbation | random | 3.8M | 3.921702014124539 | 0.7469799182493154 | 1.3887072229250794 | 0.2296470925128836 | 62 |
| 104M | Fully Frozen | Norman Perturbation | random | 7.2M | 3.7264800149972523 | 0.7682701327746247 | 1.420816461576905 | 0.21056510203332218 | 62 |
| 104M | Fully Frozen | Norman Perturbation | random | 12M | 3.858509292890057 | 0.7473774940781356 | 1.4286067679546504 | 0.22461926093192314 | 62 |
| 104M | Partially Unfrozen | Norman Perturbation | balanced | 266k | 3.1118196341255318 | 0.8249908273299993 | 1.2072555714226514 | 0.14743206520816732 | 62 |
| 104M | Partially Unfrozen | Norman Perturbation | balanced | 314k | 3.4478027971020233 | 0.8095697228364218 | 1.405716484583202 | 0.1512006405643437 | 62 |
| 104M | Partially Unfrozen | Norman Perturbation | balanced | 421k | 3.4624311166455612 | 0.8193877566716575 | 1.4629569113528944 | 0.12417748298284771 | 62 |
| 104M | Partially Unfrozen | Norman Perturbation | balanced | 647k | 3.2019108917564503 | 0.8346416971463136 | 1.3309708336658388 | 0.14379227001962644 | 62 |
| 104M | Partially Unfrozen | Norman Perturbation | balanced | 1.1M | 3.3538393057625577 | 0.82096428 | 1.3227267993274112 | 0.1377067883030213 | 62 |
| 104M | Partially Unfrozen | Norman Perturbation | balanced | 2M | 3.3893070899681796 | 0.8056701535297728 | 1.3885671499730932 | 0.17010588894807963 | 62 |
| 104M | Partially Unfrozen | Norman Perturbation | balanced | 3.8M | 3.2971974171360046 | 0.8255252172649342 | 1.2593256162155417 | 0.1430102902676957 | 62 |
| 104M | Partially Unfrozen | Norman Perturbation | balanced | 7.2M | 3.3573459917965143 | 0.7887323847752088 | 1.4326875184313241 | 0.18690047330249823 | 62 |
| 104M | Partially Unfrozen | Norman Perturbation | balanced | 12M | 3.306849288 | 0.7967504021060023 | 1.2628789151958981 | 0.17458495588726577 | 62 |
| 104M | Partially Unfrozen | Norman Perturbation | full corpus | 34M | 3.1341458480669364 | 0.8346632796749589 | 1.1532779506122077 | 0.15670365382167398 | 62 |
| 104M | Partially Unfrozen | Norman Perturbation | random | 266k | 3.2198496382828004 | 0.821341754 | 1.4262658886991377 | 0.1716172603698933 | 62 |
| 104M | Partially Unfrozen | Norman Perturbation | random | 314k | 3.3165239431157914 | 0.8157732640341541 | 1.2325313836488392 | 0.1614922488754401 | 62 |
| 104M | Partially Unfrozen | Norman Perturbation | random | 421k | 3.340220129576464 | 0.8204905529544074 | 1.495243171224673 | 0.16625327173016136 | 62 |
| 104M | Partially Unfrozen | Norman Perturbation | random | 647k | 3.1732574337162425 | 0.830180254 | 1.315581304602042 | 0.15272875895067173 | 62 |
| 104M | Partially Unfrozen | Norman Perturbation | random | 1.1M | 3.240549373147227 | 0.8370777144037921 | 1.2412018704619274 | 0.14005910098011462 | 62 |
| 104M | Partially Unfrozen | Norman Perturbation | random | 2M | 3.3115328448686125 | 0.8133012179820338 | 1.434812679 | 0.1597720481521726 | 62 |
| 104M | Partially Unfrozen | Norman Perturbation | random | 3.8M | 3.2461866521969713 | 0.8190968063927236 | 1.2153864205921427 | 0.14507593840982252 | 62 |
| 104M | Partially Unfrozen | Norman Perturbation | random | 7.2M | 3.143696329526615 | 0.8345584718836986 | 1.245365231356417 | 0.14601636620077812 | 62 |
| 104M | Partially Unfrozen | Norman Perturbation | random | 12M | 3.313303893 | 0.7975197228886054 | 1.2600499597575612 | 0.1843600026225607 | 62 |
